# MauE from Calditrichota and Thermodesulfobacteriota reveal a new pathway for disulfide bond formation in bacteria

**DOI:** 10.64898/2026.03.05.709764

**Authors:** Cassandra P. Gonzalez, Antti Moilanen, Kati Korhonen, Nguyen Pham Anh Thu, Joonatan Hiltunen, Mirva J Saaranen, Lloyd W. Ruddock

## Abstract

Disulfide bond formation is crucial to the structure and function of many proteins. It is known that there is diversity in the pathways for disulfide bond formation in bacteria and that there are gaps in our knowledge of these pathways. Using a combination of experimental and bioinformatic approaches we show that some of these gaps can be filled by a newly discovered oxidative folding pathway centered on methylamine utilization protein E (MauE). MauE has previously been associated with the methylamine utilization (MAU) gene cluster, which is involved in methylamine metabolism, in particular it is associated with the maturation of the small subunit of methylamine dehydrogenase. Here we show MauE from *Caldithrix abyssi* and *Desulfatibacillum alphaticivorans* functionally replace disulfide bond formation protein B (DsbB) in *E. coli* using two independent disulfide bond dependent assays. Furthermore, MauE is found in 14 species from 2 bacterial phyla that lack known pathways for structural disulfide bond formation, but which have proteins with structural disulfide bonds in the protein data bank. The active site for MauE was determined to be a conserved CXC motif. Using molecular docking predictions, we demonstrate that MauE is likely to interact with ubiquinone, similarly to the well characterized bacterial DsbB. We also constructed a dataset across thirty-five different phyla to demonstrate that MauE is potentially the second most common disulfide bond formation protein in bacterial disulfide bond formation pathways after DsbB. In addition, the distribution of MauE largely differs from the distribution of other MAU gene cluster markers affirming its role as a newly discovered generalist disulfide bond formation protein rather than being a specialized maturation factor for methylamine dehydrogenase. We also reveal further gaps in disulfide bond pathways, as well as species which may contain redundancies in their disulfide bond pathways.

## Introduction

Disulfide bonds are post-translational modifications that are crucial to the tertiary and quaternary structure of many proteins found throughout the tree of life. Disulfide bonds form through the oxidation of free thiols found in cysteine side chains, forming a covalent bond between them. This reaction involves the transfer of two electrons from the free thiols and is often paired with the direct (eukaryotic systems) or ultimate (some prokaryotic systems) reduction of oxygen [1,2]. During oxidative protein folding, disulfide bonds can form spontaneously depending on the cysteine pairs’ local environment and redox environment, however this process is usually very slow. Therefore, structural disulfide bonds are actively imparted through various enzymatic pathways. These pathways and their enzymatic components vary across species. In bacteria, the most well characterized oxidative pathway centers around the protein DsbB. This pathway involves DsbB and DsbA, both of which are necessary for the proper formation of disulfide bonds in the proteome of many bacterial species [3,4]. DsbB’s primary role in this pathway is as a disulfide bond formation protein, oxidizing the active site cysteines of DsbA which in turn oxidizes the thiols of a folding protein, creating a structural disulfide bond [3,4]. DsbB interacts with the quinone pool of a bacterial cell, coupling its activity to the respiratory chain of the cell [5].

The DsbB dependent pathway for disulfide bond formation was thought to be the only pathway for disulfide bond formation in the periplasm or outer membrane of bacteria until 2008 when it was demonstrated that a bacterial homolog of vitamin K epoxide reductase (VKOR) could functionally replace DsbB in DsbB *E. coli* knockout strains [6]. Through a combined approach involving computational analysis of the disulfide bonds within the proteomes of a set of bacterial species and experimental validation of VKOR as a functional disulfide bond formation protein in a new pathway for oxidative folding, Dutton *et al*. demonstrated that there is likely a diversity of pathways for the formation of disulfide bonds in bacteria. Beyond this, Dutton *et al*., demonstrated that there were gaps in the known pathways for disulfide bond formation in bacteria and highlighted some bacterial species that have either a DsbA homologue and/or structural disulfide bonds in their proteome but which lack the known disulfide bond formation proteins DsbB or VKOR. To date, as far as we are aware, none of these gaps have been filled.

MauE, is a component of the MAU gene cluster. The primary product of the MAU gene cluster is methylamine dehydrogenase (MADH), which is involved in methylamine metabolism [7]. Both MauE and MauD have previously been shown to be required for maturation of MADH, with MauD being involved in disulfide exchange with MADH allowing for the proper formation of MADH’s six disulfide bonds necessary for activity [8–10]. As MauD like DsbA contains a thioredoxin fold with a CXXC active site, most likely MauD plays the same functional role in structural disulfide bond formation in MADH as DsbA, i.e. as an intermediary protein transferring disulfide bonds from a functional homologue of DsbB or VKOR to the folding protein. Due to MauD’s involvement in the formation of disulfide bonds in MADH, and the requirement for both MauE and MauD for the maturation of MADH, we hypothesized that MauE could also be involved in disulfide bond formation, specifically that it could function as a DsbB-like disulfide bond formation protein in a new oxidative folding pathway.

In this work we demonstrate that MauE homologs from two bacterial species (*Desulfatibacillum aliphaticivorans* and *Caldithrix abyssi*) from separate phyla can functionally replace DsbB activity in *E. coli*. Furthermore, analysis of protein structures in the Protein Data Bank (PDB) reveals that proteins from 14 additional bacterial species across 2 phylum contain structural disulfide bonds, have MauE, but lack the known pathways for disulfide bond formation in bacteria DsbB or VKOR. This suggests that MauE is a widespread catalyst of structural disulfide bond formation. We then examined the prevalence and characteristics of MauE across 35 bacterial phyla and found that MauE is probably the second most common disulfide bond formation protein known so far, by both species count and distribution across bacterial phyla.

## Results and discussion

### Disulfide bond formation activity of wild type MauE

The MAU cluster has several proteins necessary to produce MADH, two of which are MauE and MauD [8]. MauD was found by Palen *et al*. to have a thioredoxin like motif (CXXC). Such motifs are found in DsbA, DsbC and DsbG and in eukaryotic catalysts of disulfide formation such as the protein disulfide isomerase (PDI) family [4,11–13]. Hence, MauD having this motif was taken as an indicator of its role in the proper formation of disulfide bonds in the MADH complex. More recently Alvarez *et al*. investigated MauE and found it to potentially be a reductase [14]. However, it is unclear how it could function as a reductase as it lacks many of the features of known components of the reducing pathways such as DsbD in bacteria [15,16] and it was shown to be more sensitive to the reductant dithiothreitol (DTT) implying a role in oxidation rather than reduction. As it has been reported that unknown DsbB-like disulfide bond formation proteins must exist [6], and MauE is found in some species (e.g., *Solibacter usitatus, Geobacter sulfurreducens*) which represented known gaps in disulfide bond formation pathways amongst bacteria published by Dutton *et al.,* in 2008, we hypothesized that MauE may be a disulfide bond formation protein.

Both known bacterial disulfide bond formation proteins, DsbB and VKOR have a core four transmembrane helix bundle structure with a pair of cysteine residues forming their active site [17,18]. To examine if MauE could potentially also be a disulfide bond formation protein, we first compared the predicted structure, topology and presence of potential active site cysteines of two MauE homologues, Uniprot-ID: B8FLW7 from *Desulfatibacillum aliphaticivorans* and H1XRV2 from *Caldithrix abyssi*. This was then compared with those of DsbB and VKOR. MauE was predicted to have a similar structure to that of DsbB and VKOR by AlphaFold2 [19,20], with a topology placing a pair of cysteines arranged in a CXC motif in the periplasm as predicted by TOPCONS [21] (Fig 1, S1 Fig). This supports the concept that MauE may play a similar functional role as a disulfide bond formation protein.

**Fig 1.**
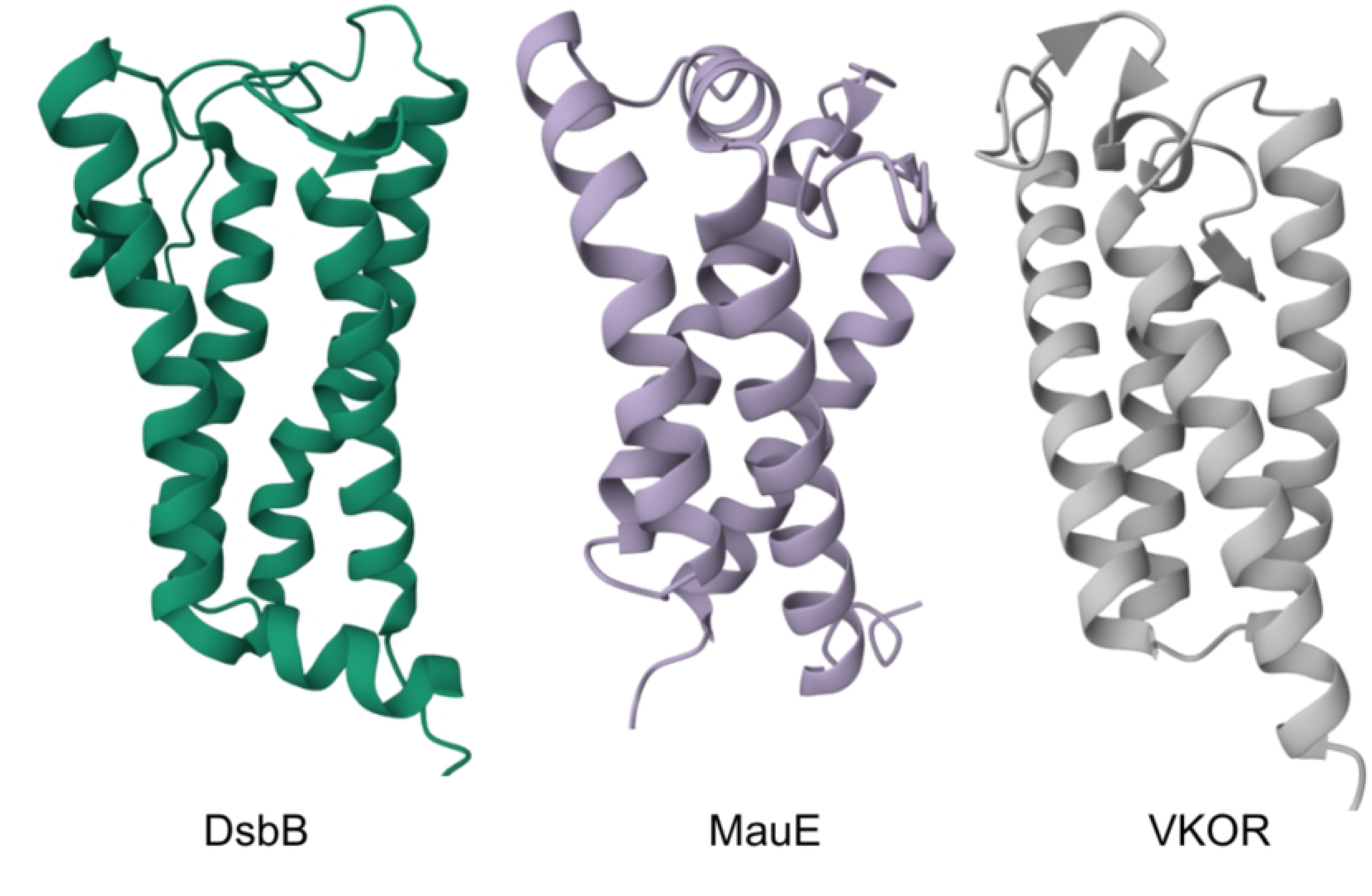
Comparison of the structures of transmembrane proteins involved in disulfide bond formation. *Escherichia coli* DsbB [22] (PDB: 2K74), and *Caldithrix abyssi* MauE (Uniprot ID: H1XRV2) and *Vermiphilus pyriformis* VKOR (Uniprot ID: A0A0D2GNY3) from AlphaFold-2 prediction [19].

The disulfide bond formation proteins in oxidative protein folding usually do not interact directly with the folding protein, but rather they oxidize a thioredoxin superfamily member, for example DsbA or PDI [3,23]. In eukaryotes and prokaryotes these two functions can exist as a fusion protein, for example 9.3% of all proteins annotated with a VKOR InterPro motif exist in a fusion with a thioredoxin superfamily member [18] (S2 Fig). Examination of all InterPro MauE protein entries shows a similar level of existence of fusions with a thioredoxin superfamily member (8.2%; S2 Fig). This supports the hypothesis that MauE is similarly a disulfide bond formation protein and suggests that MauE may work via an intermediary protein that is a member of the thioredoxin superfamily e.g. DsbA or MauD. In the InterPro database there are more proteins annotated as MauE (19,775) than VKOR (16,690) and only slightly less than DsbB (21,378). This suggests that MauE could be the second most prevalent DsbB-like disulfide bond formation protein in bacterial species.

To evaluate the ability of MauE to functionally replace DsbB, two distinct assays previously used in the field to examine the replacement of DsbB activity by VKOR were undertaken [6]. MauE from *Desulfatibacillum aliphaticivorans* (B8FLW7) and *Caldithrix abyssi* (HX1RV2) were first assessed through a bacterial motility assay. In *E. coli* motility requires correct folding of FlgI, a periplasmic protein associated with the P-ring of the flagellum that contains one intramolecular disulfide bond in its native state. This disulfide bond is required for motility [24]. When either B8FLW7 or HX1RV2 were introduced into Δ*dsbB E. coli*, motility was restored and was comparable to the motility of the K12 wildtype *E. coli* positive control (Fig 2A).

**Fig 2.**
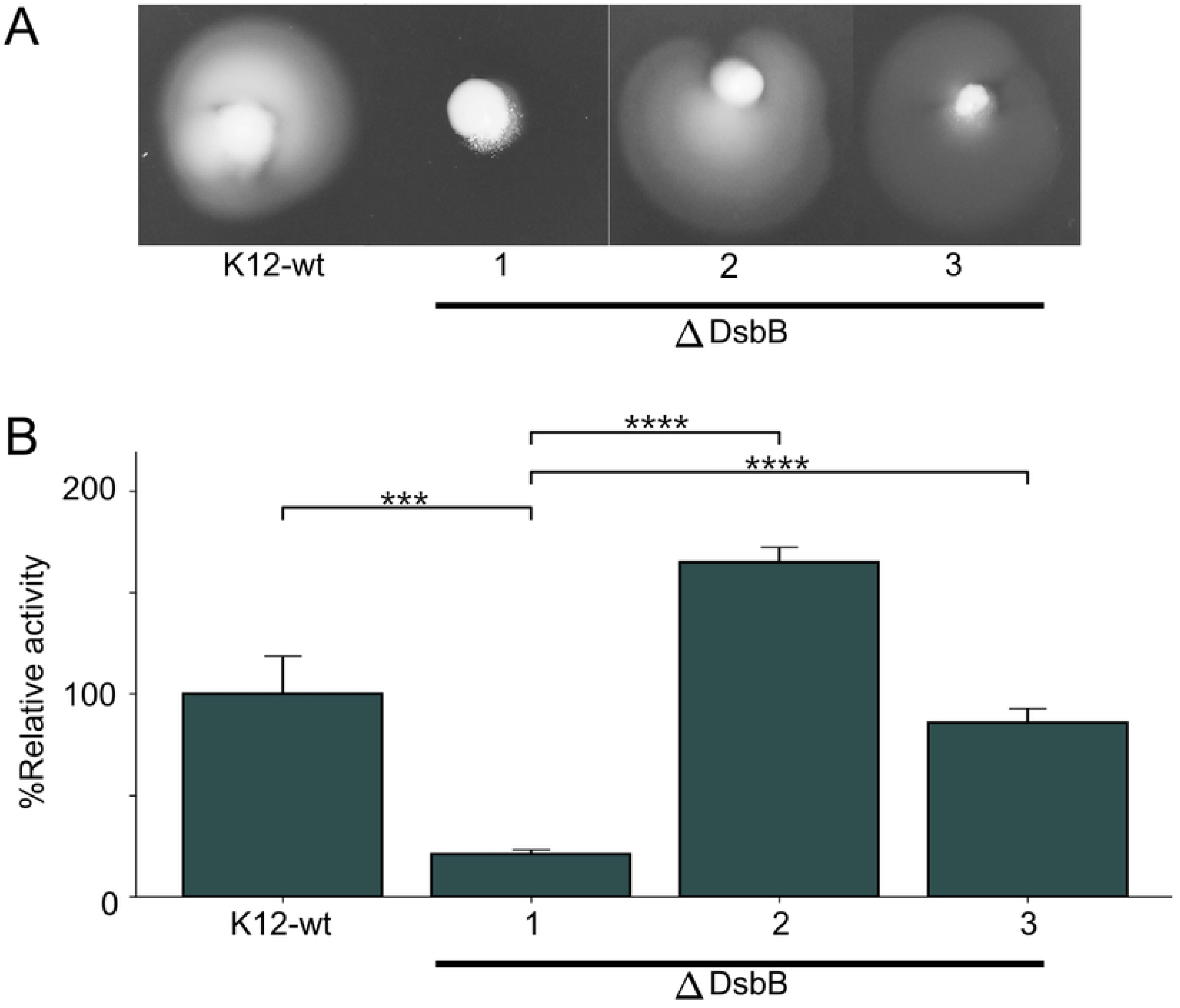
Disulfide bond formation activity assays for wild-type MauE. (A) B8FLW7 in Δ*dsbB* K12 *E. coli* (2) and H1XRV2 in Δ*dsbB* K12 *E. coli* (3) showed motility comparable to that of the K12 wildtype *E. coli* and surpassing that of Δ*dsbB* K12 *E. coli* (1). Representative motility tests are shown. (B) The relative activity of endogenous PhoA in both B8FLW7 in Δ*dsbB* K12 *E. coli* (2) and in H1XRV2 in Δ*dsbB* K12 *E. coli* (3) significantly exceeded that of the Δ*dsbB* K12 *E. coli* negative control (1) and was comparable to or greater than that of the K12 wildtype *E. coli*. The graph shows the mean and standard deviation of the data. Four asterisks indicate p-values < 0.0001 and three indicate p-values < 0.001. P-values were calculated using a two tailed two sample unequal variance T-test for each group compared to Δ*dsbB* K12 *E. coli*. N = 6 for each group.

The same two MauE were then assessed for their ability to properly impart disulfide bonds in the periplasmic protein alkaline phosphatase (PhoA) through a colorimetric activity assay. PhoA has two disulfide bonds that play roles in activity and stability, and without these disulfide bonds activity is drastically reduced [25]. When either B8FLW7 or HX1RV2 were introduced into Δ*dsbB E. coli*, PhoA activity was rescued to levels equivalent to the wildtype K12 *E. coli* positive control or higher (Fig 2B).

These two independent assays show that MauE from two separate bacterial phyla can functionally replace DsbB in a Δ*dsbB* knockout strain of *E. coli*, i.e. that MauE can function as a DsbB-like disulfide bond formation protein.

### Identification of the active site of MauE

Both DsbB and VKOR contain redox active cysteines in their active sites [18,26], and we hypothesized that this would be the same for MauE. For both B8FLW7 and HX1RV2, there are only two cysteines present in their protein sequences. These form a CGC motif as part of a disordered loop between the third and fourth transmembrane helices (S1 Fig). To determine experimentally if the CXC motif forms part of the active site, either the N-terminal cysteine (referred to hereafter as C1), the C-terminal cysteine (referred to hereafter as C2), or both cysteines were mutated to alanine. The mutants of MauE were then examined for their ability to make disulfide bonds in Δ*dsbB E. coli*.

While expression of wildtype B8FLW7 and H1XRV2 restored motility in a Δ*dsbB* strain, none of the CXC site mutants, either single or double, restored motility (Fig 3A). Similarly, both the single and double mutants failed to restore PhoA activity, while wild-type B8FLW7 and H1XRV2 did (Fig 3B). These results implicate the CXC motif in MauE as being required for activity. Most likely these form the active site, as per the redox active cysteines in the active sites of DsbB and VKOR [18,26].

**Fig 3.**
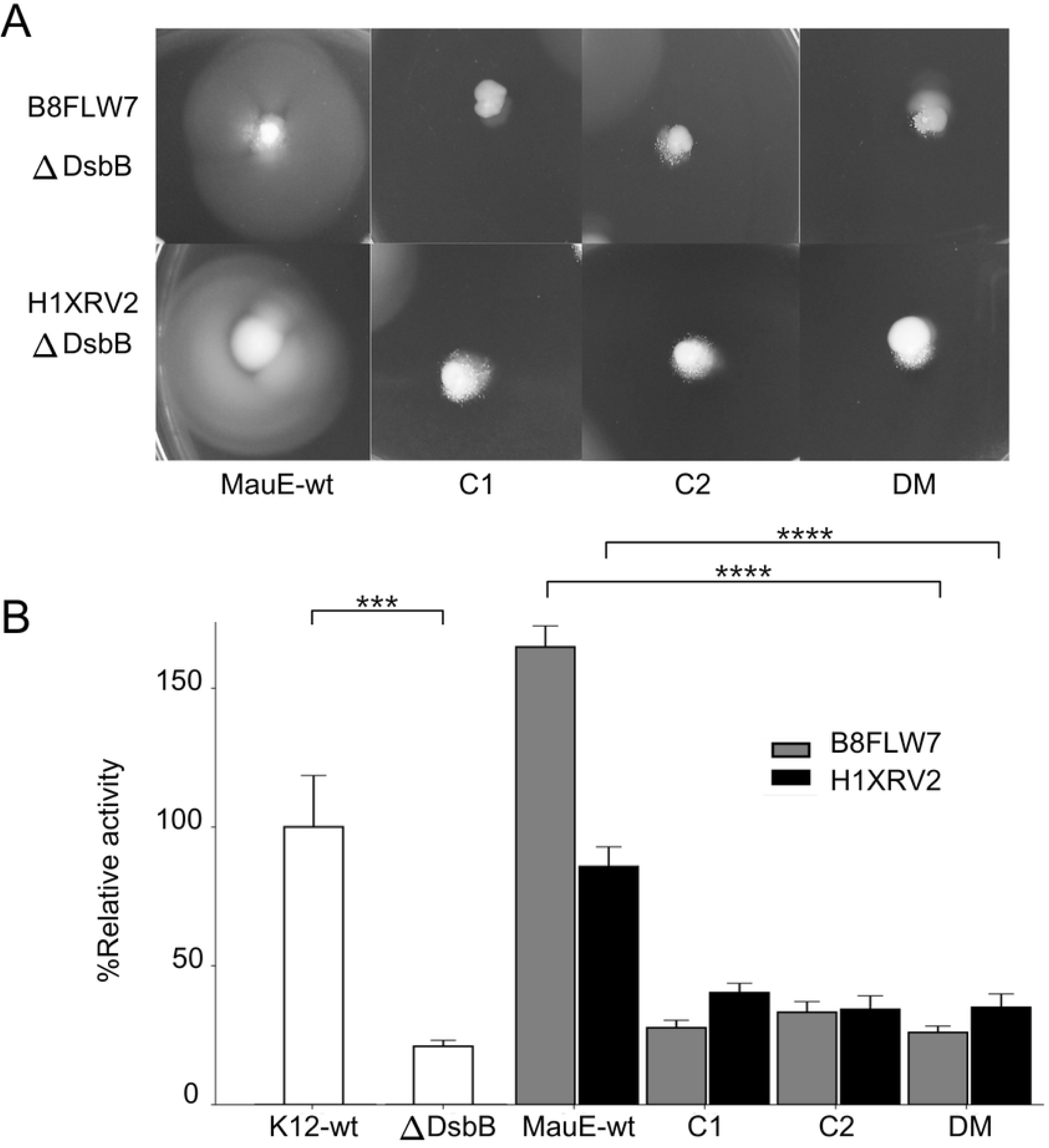
Disulfide bond formation activity assays for MauE with the CXC motif mutated. (A) The motility of B8FLW7 in Δ*dsbB* K12 *E. coli* and H1XRV2 in Δ*dsbB* K12 *E. coli* was severely disrupted when either cysteine (C1 or C2) or both cysteines (DM) in the CXC active site were mutated to alanine. Representative motility tests are shown. (B) The relative activity of endogenous PhoA in B8FLW7 in Δ*dsbB* K12 *E. coli* and H1XRV2 in Δ*dsbB* K12 *E. coli* was severely disrupted when either cysteine in the CXC active site was mutated to alanine. The graph shows the mean and standard deviation of the data (n = 6 for each group). Four asterisks indicate p-values < 0.0001 and three indicate p-values < 0.001. The p-values were calculated comparing wildtype B8FLW7 and H1XRV2 to their respective double cysteine to alanine mutants using a two tailed two sample T-test. B8FLW7 used an unequal variance T-test, and H1XRV2 used an equal variance T-test as determined by performing an F-test on their respective variances in comparison to the double mutant.

The location of the CXC motif in a disordered loop on the periplasmic side of MauE is reminiscent of the CXXC active site motif of DsbB. This loop in DsbB can adopt multiple conformations and can interact with ubiquinone buried within the transmembrane helices, coupling disulfide bond formation to the respiratory chain of the bacteria [1,26]. A similar dynamic loop containing a CXXC active site is suspected to shuttle electrons between a quinone acceptor bound to VKOR and VKOR’s thioredoxin like binding partner [18]. To investigate the possibility that the CXC motif of MauE could be making a similar interaction with ubiquinone, docking simulations using GNINA [27] (molecular docking software fork of Autodock VINA and smina) were performed. GNINA predicted a favorable interaction with ubiquinone-8, placing ubiquinone-8 within the center of the transmembrane helices of DsbB with the quinone group positioned near the CXXC active site of DsbB as expected (S3A Fig). Both MauE were also predicted to favorably interact with ubiquinone, with GNINA placing ubiquinone-8 between the transmembrane helices of both MauE in a manner similar to the docked position of ubiquinone-8 in DsbB (S3B-C Fig). As a negative control, a decoy molecule, palmitic acid, was docked to all three proteins. GNINA scores for DsbB, B8FLW7 and H1XRV2 docking palmitic acid were overall lower with the docked position often occurring outside the transmembrane helices (S3D Fig). GNINA does not allow major protein backbone movements during docking which makes movements of the CXC disorder loop of MauE difficult to predict. To determine what potential protein backbone movements of the CXC loop might look like, the structure of the ubiquinone-8 bound state of both MauE was predicted with the AlphaFold-3 replication model Protenix [28]. The bound state structure predictions highlighted the coordination of the quinone head by glutamic acid and lysine side chains along with the movement of the CXC loop into a favorable position for interaction with ubiquinone-8 for both MauE (Fig 4). These docking predictions indicate that both B8FLW7 and H1XRV2 could interact with ubiquinone in a comparable manner to DsbB, suggesting that the two proteins share a comparable route for disulfide formation.

**Fig 4.**
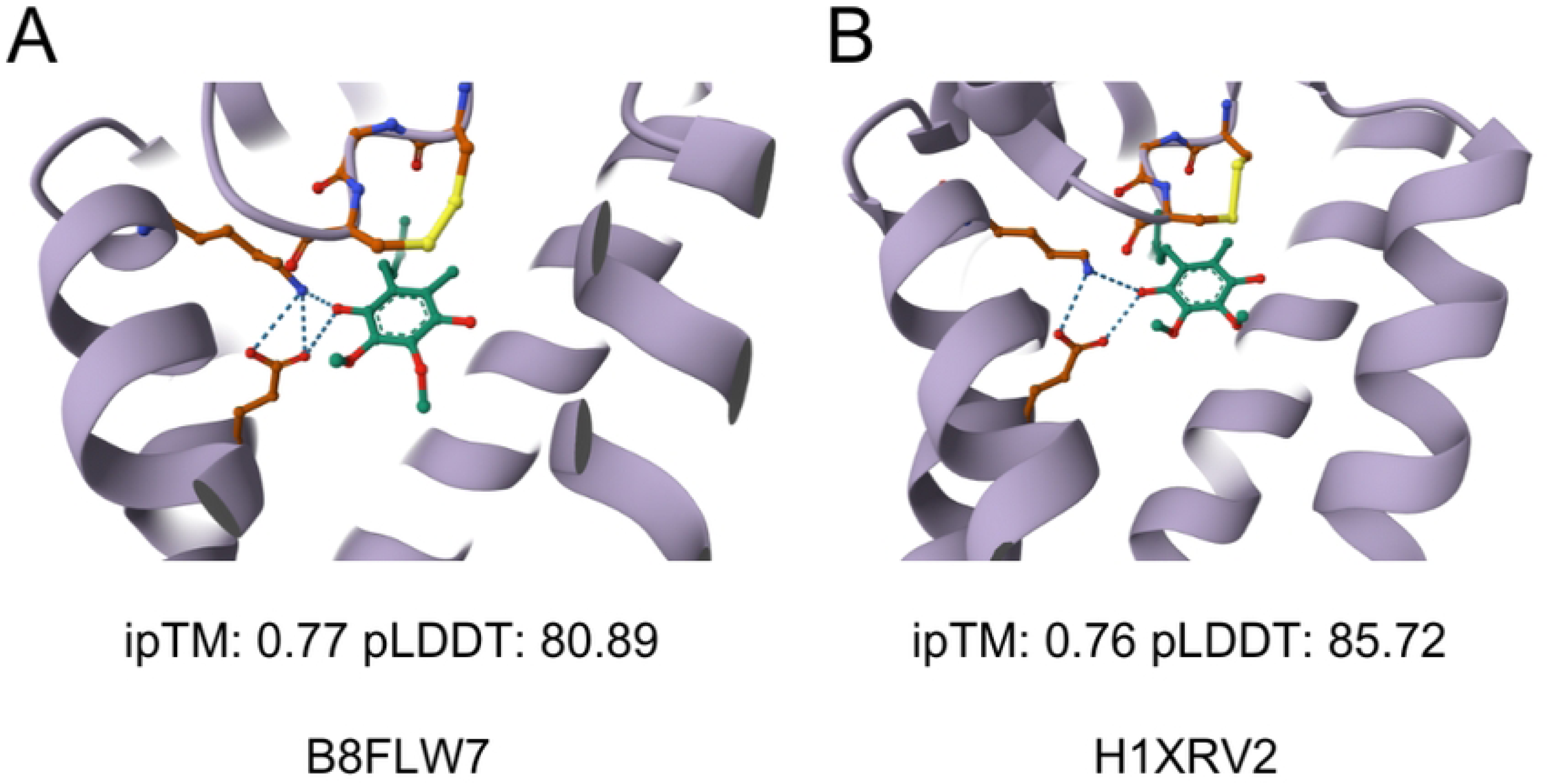
Docking of ubiquinone-8 with MauE. (A-B) The predicted bound state of ubiquinone-8 (green) with B8FLW7 (A) and H1XRV2 (B). The quinone head is positioned within the space between the transmembrane helices in both MauE, allowing juxtaposition with the CXC motif in the disordered loop and interactions with a conserved glutamic acid and lysine (orange). The listed ipTM and pLDDT values were taken from the top ranked predictions output from Protenix.

### Phylogenetic analysis of MauE utilization

*In vivo* assays show that MauE from two different phyla are disulfide bond formation proteins capable of functionally replacing DsbB and that they contain a CXC active site. This raises several immediate questions. What is the conservation of amino acid identity near the active site of MauE? Is MauE present only when the MAU cluster is present? How widely is MauE distributed across bacterial phyla and across bacterial species? To investigate these issues, a phylogenetic analysis was performed encompassing 20,704 bacterial species across thirty-five different phyla.

The total initial MauE protein entries within the data set exceeded 19,000, however only 15,413 contained a CXC active site and met the quality cut off for analysis (see methods). The amino acid sequences for the 15,413 MauE protein entries containing a CXC active site were used to analyze the amino acid variation around the CXC active site. It was found that the X position was often a glycine residue, and that the CXC motif was followed by a highly conserved phenylalanine and glycine (Fig 5A). A smaller subset of 4,000 non-redundant MauE protein entries were then randomly selected from the entire set of 15,413 MauE to find conserved amino acid identities through the entirety of the MauE sequences. A highly conserved lysine and glutamic acid were found at positions on the first and second transmembrane helices of H1XRV2 and B8FLW7 near the region ubiquinone-8 was predicted to bind for both H1XRV2 and B8FLW7 by GNINA and Protenix (S4 Fig). In DsbB it is thought that two conserved phenylalanine residues (Phe-32 and Phe-106) are involved in binding ubiquinone in a manner like that of NADH-quinone acceptor oxidoreductase [26]. Given the high degree of conservation of the phenylalanine after the CXC motif of MauE, and the high degree of conservation of the lysine and glutamic acid near the region where ubiquinone is predicted to bind to H1XRV2 and B8FLW7, it is possible that these three amino acids aid in ubiquinone binding during the catalytic cycle of MauE.

**Fig 5.**
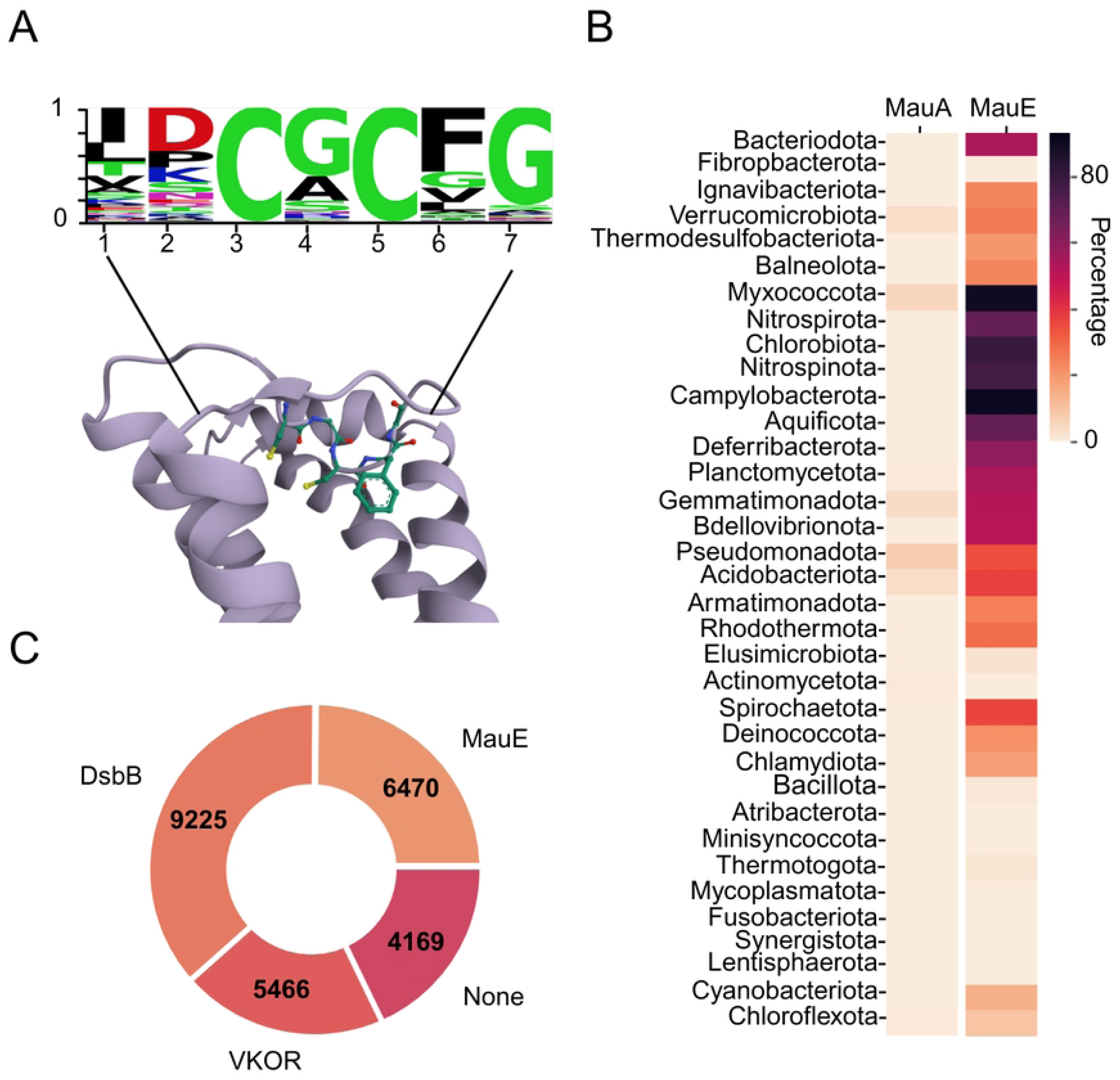
The distribution of MauE across bacterial phyla, and its active site amino acid distribution. (A) The amino acid sequence distribution of all MauE homologs containing a CXC motif in our dataset. The X site is dominated by the smallest amino acids glycine or alanine and a highly conserved phenylalanine and glycine follow the CXC motif. (B) The distributions of MauA and MauE across the 35 analyzed bacterial phyla, showing no similarity in distributions of the two. (C) The number of species in the dataset of 20,704 species containing MauE, DsbB, and VKOR or none of the three. MauE is the second most prevalent of the three in our dataset. The sum of these numbers exceeds the total number of bacterial species with these IPRs due to some species having multiple members.

We then examined whether the presence of MauE correlated to the presence of the MAU gene cluster in which it was first identified. For this we used the marker protein MauA. MauA is the light chain subunit of MADH which alongside MauB makes up the tetrameric complex of MADH [29]. MauE was originally identified as a protein involved in the maturation of MauA [8]. While 6,470 species in our dataset contained MauE only 659 of these also contained MauA, i.e. only 10.2% of species that have MauE also contain MauA. Furthermore, examination of the phyla distribution of MauE and MauA reveals that MauE is present in many phyla in which MauA is not (Fig 5B). This implies that MauE is not strictly tied to the maturation of MADH in the MAU cluster, but rather that it is a generalist maturation factor. This is consistent with a role in oxidative folding. The MAU cluster also contains MauG which is also involved in the maturation of MauA, via its involvement in the synthesis of tryptophan tryptophylquinone [30]. Phylogenetic analysis revealed that MauG is also much more widespread than MauA and that its distribution does not match that of MauE (S5 Fig).

The dataset was then used to determine the per phylum usage of the three disulfide bond formation proteins, MauE, DsbB and VKOR. Consistent with the relative abundance in the InterPro database, MauE was found to be the second most prevalent pathway for disulfide formation after DsbB by bacterial species count (Fig 5C). Furthermore, MauE showed a wide breadth of distribution across bacterial phylum, with a large cluster of bacterial phyla with high percentages of bacterial species which contain MauE (S6 Fig). Consistent with the smaller dataset from Dutton *et al.* [6] and analogous to the situation in eukaryotes [31,32], a significant number of species were found to contain representatives from multiple different classes of disulfide bond formation proteins (S6 Fig).

Since biochemical data on two MauE combined with IPR analysis suggests that MauE is a prevalent pathway for disulfide bond formation across a large number of bacterial species, we then considered if other supporting data was available from other databases. The protein data bank (PDB) contains over 245,000 protein structures. Of these circa 10,035 are annotated as bacterial protein structures containing at least one disulfide bond. These were then examined on a species-specific level, with additional filtering to ensure that the disulfide bond observed was likely to be structural rather than functional (see methods). Examination of proteins from species which lack both DsbB and VKOR but which contain MauE revealed that 21 periplasmic or secreted proteins from 14 species from 2 phylum in PDB are reported to have structural disulfide bonds (Table 1). This supports the concept that MauE is a novel widely spread pathway for disulfide formation.

**Table 1.**
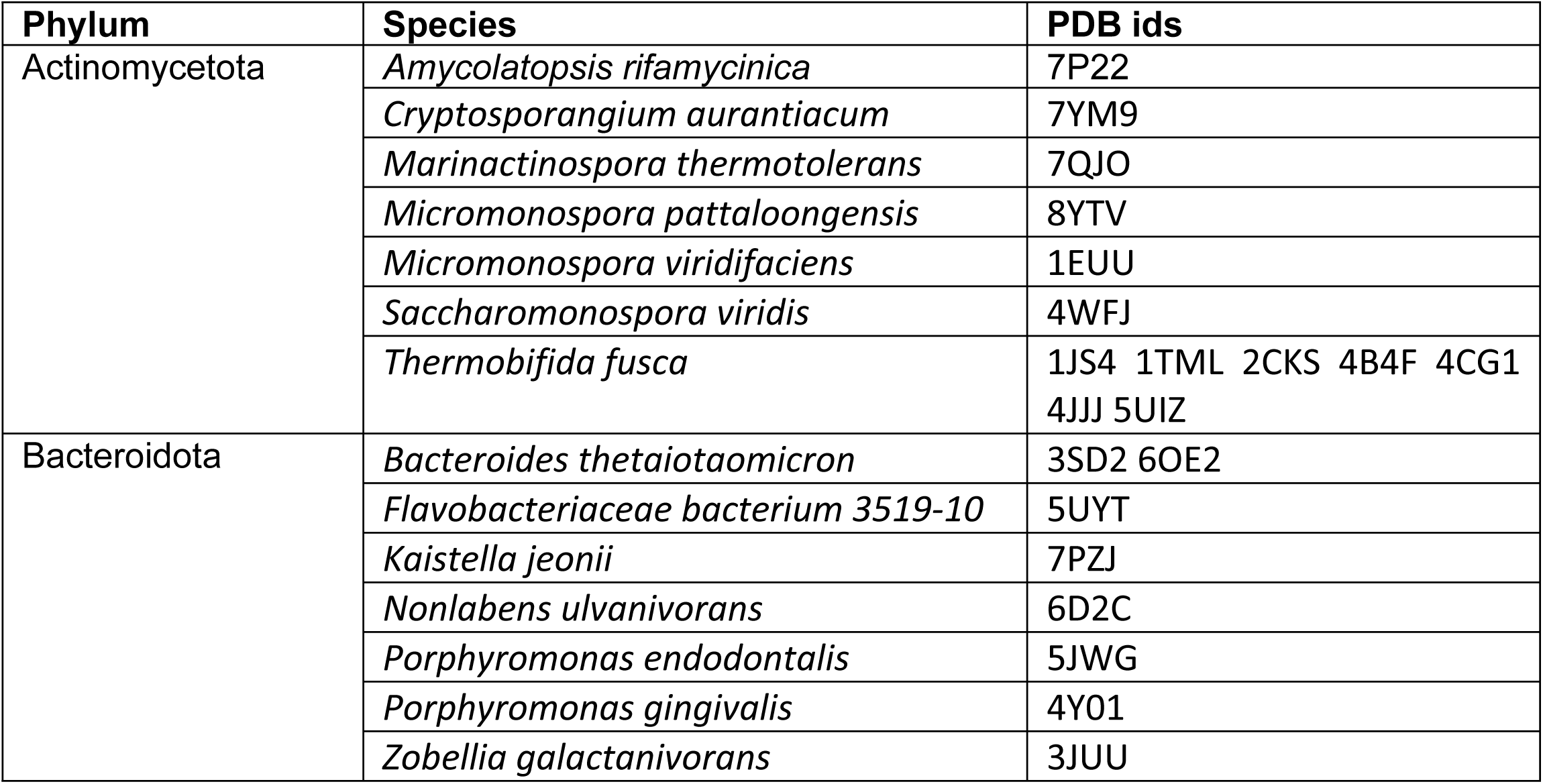
PDB entries of proteins that are translocated from the cytoplasm to the periplasm, membrane or are secreted, that have at least one structural disulfide and are found in species that have MauE but lack DsbB or VKOR.

With the discovery of MauE as a generalist disulfide bond formation protein, there is the question of whether there are as yet unidentified pathways for disulfide bond formation. In our dataset we found 4,169 species (20.1% of the data set) which lack DsbB or VKOR or MauE. Some of these bacterial species may not contain proteins with structural disulfide bonds, but it is likely that many do, and at least some of the gaps in disulfide bond pathways identified by Dutton *et al*. [6] are also represented in this category of our dataset. We then undertook the same PDB based analysis for these species as for MauE containing species. Again, despite general underrepresentation of these species in the PDB, 22 proteins from 13 species in 6 phyla, were identified as containing structural disulfide bonds from species which lack DsbB or VKOR or MauE (Table 2). This indicates that there are more pathways for oxidative folding still to uncover. It is not unreasonable to think that some of these as yet unidentified pathways could be of interest in bio-industrial applications or as targets for antibiotic drugs. It would be in our interest to uncover more of these pathways if not only for the benefit of deepening our understanding of protein folding.

**Table 2.** PDB entries of proteins that are translocated from the cytoplasm to the periplasm, membrane or are secreted, that have at least one structural disulfide and are found in species that lack DsbB, VKOR and MauE.

| Phylum | Species | PDB IDs |
| --- | --- | --- |
| Aquificota | *Aquifex aeolicus* | 2DI4 3HYV |
|  | *Sulfurihydrogenibium azorense* | 4X5S |
|  | *Thermovibrio ammonificans* | 4C3T |
| Bacillota | *Acetivibrio thermocellus* | 5GXX |
|  | *Clostridium perfringens* | 1UPS |
|  | *Staphylococcus aureus* | 1CQV 1DYQ 1D5M 1ENF 1JCK<br>1XXG 3OWE 4UDU |
|  | *Streptococcus pyogenes* | 1B1Z 1BXT |
| Campylobacterota | *Helicobacter hepaticus* | 7S9Z |
|  | *Wolinella succinogenes* | 3KN3 |
| Chlorobiota | *Chlorobaculum tepidum* | 8HN2 |
| Planctomycetota | *Candidatus Jettenia caeni* | 5ZL1 |
|  | *Candidatus Kuenenia stuttgartensis* | 6HIF |
| Spirochaetota | *Borrelia burgdorferi* | 2LQU |

## Materials and Methods

### Molecular Biology

The Δ*endA* strain XL1-Blue (TetR) was used for all cloning and plasmid isolation steps. Synthetic genes codon optimized for *E. coli* K12, encoding H1XRV2 and B8FLW7 were ordered from GenScript with flanking NdeI and BamHI sites. The genes were cloned into pMJS163 (S7 Fig), a modified pET23a-based vector containing AmpR and the T7 promoter replaced with a pTAC promoter as described earlier [33]. Plasmid DNA was isolated using an E.Z.N.A Plasmid DNA Mini Kit 1 (Omega Bio-tek). Mutants were created using the Agilent QuickChange Site-Directed Mutagenesis Kit (Agilent Technologies), using primers shown in the S1 Table. The DNA sequences for H1XRV2 and B8FLW7 were codon optimized using the GenScript codon optimization tool and are listed in the S2 Table. Both wildtype and mutant plasmid vectors were sequenced prior to use.

### Motility Assay

Validated plasmids carrying the gene of interest were heat shock transformed into Δ*dsbB* K12 *E. coli* (JW5182, Keio collection) for activity assays and selected for with LB medium containing 100µg/ml ampicillin and 30 µg/ml kanamycin. The parent strain *E*. *coli* K-12 strain BW25113 served as the wildtype control [34,35]. Motility assay plates were created using 0.3 % agar in LB medium with 100 µg/ml ampicillin and 30 µg/ml kanamycin. Motility plates for H1XRV2 had 5 µM Isopropyl β-D-1-thiogalactopyranoside (IPTG), and motility plates for B8FLW7 had 10 µM IPTG, to induce appropriate levels of these transmembrane proteins. These IPTG concentrations were determined empirically. Freshly transformed Δ*dsbB* K12 *E.* coli or control strains were spotted onto motility plates and incubated at 30°C for 24 hours before visual assessment of motility. Motility plates were repeated three times for each group on two separate days for an N of 6 for all groups including the controls.

### Alkaline Phosphatase activity assay

The endogenous PhoA activity assay was performed as described previously [36]. H1XRV2 in Δ*dsbB* K12 *E. coli* or B8FLW7 in Δ*dsbB* K12 *E. coli* again received 5 µM and 10 µM IPTG respectively with added 100µg/ml ampicillin and 50µg/ml kanamycin. The parent strain *E*. *coli* K-12 BW25113 and Δ*dsbB* K12 *E. coli* JW5182 served as positive and negative controls respectively [34,35]. The controls, H1XRV2 in Δ*dsbB* K12 *E. coli* and B8FLW7in Δ*dsbB* K12 *E. coli,* were run in triplicate starting from three separate inoculating colonies each. This assay was performed twice for a total N of 6 for each group. All cultures were grown in sterile 50 ml centrifuge tubes with the lids sealed to disrupt gas exchange and allow for more anaerobic conditions and thus prevent background oxidation. Activity was measured by monitoring ΔA_410_ over the course of 1 hour at 25°C in a 96 well plate using a Tecan M1000 plate reader (Tecan). This was repeated for the H1XRV2 and B8FLW7 single and double mutants.

### Phylogenetic analysis

Initial datasets for phylogenetic analysis were constructed from the InterPro database [37] which also draws upon the UniProt database [38]. Data was acquired using the InterPro API using the NCBI taxonomic ID of 2 as a filter during the InterPro API pull, which corresponds to bacteria. The data set was constructed by using the IPR codes found in the S3 Table, which corresponded to protein entries for DsbB, VKOR, MauE, MauA, MauG, Bacterial Signal Peptidase, Bacterial RNA polymerase, and bacterial large ribosomal subunit. The later three were chosen to capture the majority of species contained within the bacterial entries in the InterPro database and could thus capture the “none” category. Additionally, these three highly conserved bacterial proteins were used to determine the quality of the data underlying the InterPro entries for each bacterial species, with anything not containing all three not meeting the quality requirements and being removed. All associated meta data contained within the InterPro database, including NCBI taxonomic IDs, amino acid sequence, sequence length, Source organism, database accession number, and AlphaFold database structural predictions were acquired during the API pull. Python 3.8.8 was used to parse the resulting data utilizing the Pandas, NumPy, Matplotlib and Biopython packages for all subsequent analysis [39–43]. The constructed dataset was then aggregated at the species level and then grouped by phyla. Scientific names corresponding to NCBI taxonomic IDs were acquired using the NCBI Entrez API via the Entrez system in Biopython as well as the full catalogue of information on taxonomic IDs available for download on the NCBI webpage [44]. All MauE entries that did not contain a CXC active site motif were removed. The CXC motif logo plot was created by taking the XXCXCXX positions of every MauE that had a single CXC motif and contained all three quality markers. Any MauE with CXC positions at either terminus were dropped leaving the final considered entries totaling at 15,413. These were processed using Biopython’s AlignIO module and the amino acid distribution was calculated using pandas before plotting using the Logomaker python package. The full MauE amino acid sequence logo plot was created using 4,000 randomly sampled non-redundant MauE entries from the 15,413 MauE. The random sampling was done using Random and Pandas Python packages to generate 4,000 random numbers in the range of the dataset index numbers, and checking the random samples for repeats to ensure the set had no redundant entries, before pulling the sequences associated with the dataset index. These were then aligned using Clustal omega. Amino acid positions that were not in at least 50% of the 4,000 entries were dropped before the amino acid distributions were calculated and plotted in the same manner as the CXC motif. The phylum level analysis was further filtered by removing candidate phylum, and phylum with less than 10 species entries leaving 35 phyla in the final analysis. This was repeated for the phylum level analysis of the MAU gene distributions. The code used for raw data acquisition, metadata acquisition, data annotation, and iterative filtering is provided via Github, alongside the final data tables used in analysis.

### Analysis of PDB structures that contain disulfide bonds

PDB entries of bacterial species containing at least one disulfide bond were collected using the search API tools from the PDB and Uniprot databases, including metadata on taxonomy, sequence and uniprot entries. These were then cross correlated against the final dataset used in the phylogenetic analysis to isolate species that either had MauE but no DsbB or VKOR or which lacked all three. These subset of proteins were then assessed with TOPCONS-2 [21] to identify proteins that contained transmembrane helices, or signal peptides indicating their translocation from the cytoplasm to the periplasmic space, association with the membrane or secretion. The remaining proteins were then manually filtered to remove proteins that contained only previously reported catalytic disulfide bonds e.g. DsbA, as well as proteins in which the only disulfides reported involved engineered in cysteine residues. The structures of the remaining proteins were visually analyzed and proteins that only contained disulfide bonds which appeared functional rather than structural e.g. surface exposed CXXC motifs were discarded.

### Molecular docking predictions

PDB files associated with B8FLW7 and H1XRV2 from the AlphaFold structural database were used for docking [20]. The structure of DsbB was taken from the RCSB entry 2K74 [22]. The structure of DsbB from 2K74 was prepared for docking by removing waters and any associated ligands in the structure using PyMOL 2.0 (Schrödinger). The ligands used for docking were palmitic acid and ubiquinone-8 taken from the RSCB database with the ligand downloader tool [45]. Both ligands were used for docking as.SDF files. To perform the docking we used GNINA 1.3, as it was the highest performing machine learning based tool [46] which allowed the use of both ligands. The run settings used for GNINA is available in supplementary data alongside the resulting score files.

## Acknowledgments

The use of the facilities and expertise of the Biocenter Oulu core facilities, part of Biocenter Finland, is gratefully acknowledged.

## Data availability

All data used in this work is available for download at: https://doi.org/10.5281/zenodo.18197764.

All code used in analysis is available for download at: https://doi.org/10.5281/zenodo.18199468

## Abbreviations

Dsb: disulfide bond formation protein
GNINA: molecular docking software fork of Autodock VINA and smina
IPTG: Isopropyl β-D-1-thiogalactopyranoside
MADH: methylamine dehydrogenase
MAU: methylamine utilization gene cluster
PDB: Protein Data Bank
PDI: protein disulfide isomerase
VKOR: vitamin-K epoxide reductase.

---

**S1 Fig. Structural and topological analyses of MauE.** (A) The AlphaFold-2 predicted structures of B8FLW7 (*Desulfatibacillum aliphaticivorans*) and H1XRV2 (*Caldithrix abyssi*) colored according to pLDDT confidence score with a darker blue representing higher confidence and yellow to red representing medium to low confidence [19]. (B) The respective sequences and TOPCONS predicted topologies about the periplasmic membrane of B8FLW7 and H1XRV2. Both MauE are 4 helical transmembrane proteins with the active site CXC motif between the third and fourth helix facing the periplasm. TOPCONS represents periplasmic facing regions as “o”, cytoplasmic facing regions as “i”, and membrane spanning regions as “M”.

**S2 Fig. Analysis of the existence of fusion proteins in the InterPro database for MauE, DsbB and VKOR.** Proteins annotated as containing MauE, DsbB or VKOR were split into three categories i) proteins not annotated as being in a fusion with any other IPR; ii) proteins annotated as being in a fusion with a thioredoxin-superfamily member (e.g. thioredoxin, glutaredoxin, DsbA etc) and no other IPR; iii) proteins annotated as being in a fusion with a non-thioredoxin superfamily member.

**S3 Fig. Docking of ubiquinone-8 and a palmitic acid decoy to DsbB and MauE.** (A) The predicted docking position of ubiquinone-8 within DsbB. (B-C) The docked positions of ubiquinone-8 in both MauE (B8FLW7 and H1XRV2). All three predictions place the ubiquinone head within the transmembrane helices of each protein near the periplasmic facing region where the active sites reside. (D) The predicted affinities of DsbB, B8FLW7 and H1XRV2 for ubiquinone-8 and the decoy molecule palmitic acid. All three have a high predicted affinity for ubiquinone-8 and a lower predicted affinity for palmitic acid.

**S4 Fig. The conserved amino acids of MauE.** (A) The amino acid distributions along the full length of MauE shown as a logo plot. In addition to conservation around the CXC active site motif there are a highly conserved lysine and glutamic acid highlighted by the black triangles above the amino acid positions. (B) The highly conserved lysine and glutamic acid positions highlighted on the predicted structure of H1XRV2 and B8FLW7.

**S5 Fig. Clustered heat map of three genes from the MAU gene cluster (MauA, MauE and MauG) depicting their distribution across the thirty-five bacterial phyla in this dataset.** MauA is the light chain of the MADH protein complex [23]. MauG is involved in the formation of the tryptophan tryptophylquinone cofactor of MauA [18]. The distribution of MauA does not correlate directly to the distribution of MauE or MauG, with both MauE and MauG showing a much wider distribution. The distribution of MauE and MauG also do not correlate directly. These results are consistent with both MauE and MauG being generalists, rather than specific enzymes required for the maturation of just MauA.

**S6 Fig. Clustered heat map of MauE, VKOR and DsbB depicting their distribution across the thirty-five bacterial phyla in this dataset.** The percentage usage of each of the three disulfide bond formation proteins, some combination of the three or none of the three per bacterial phyla by species counts. The color scale of the heat map represents the percentage of species in the phyla that use the respective category of enzyme, with lighter colors representing a higher percentage of species. The heatmap is clustered to group phyla with similar usages closer together. The clustering reveals a group of eight bacterial phyla that have higher percentages of species that only use MauE and contain neither DsbB nor VKOR. This cluster is the second largest cluster. A small population of bacterial species using all three enzymes is present in several phyla such as *Bdellovibrionota*.

**S7 Fig. Plasmid map for pMJS163**

**S1 Table. DNA primers used in this work.** The table lists primers used for site directed mutagenesis for the C1 and C2 mutants of H1XRV2 and B8FLW7. The names denote the amino acid position of the Cysteine which was mutated to an Alanine. The table also lists the sequencing primers used to assess the resulting mutants using sanger sequencing.

**S2 Table. DNA sequences of both MauE homologs.** The DNA sequences of B8FLW7 and H1XRV2 codon optimized for *E. coli* by GenScript.

**S3 Table. InterPro codes used for dataset construction.** The InterPro IPR codes used for acquiring all the InterPro entries that make up the dataset used in this work. IPR047873, IPR010243 and IPR036286 were chosen because they are highly distributed across bacterial species and can thus capture a large sample of bacterial species contained within the InterPro database, including those which do not have any instance of MauE, DsbB, VKOR or MauA. Additionally, they serve as quality markers for the underlying sequencing, with quality entries containing all three for the respective species.

